# DNA hypomethylation promotes the expression of CASPASE-4 which exacerbates neuroinflammation and amyloid-β deposition in Alzheimer’s disease The Ohio State University College of Medicine

**DOI:** 10.1101/2023.08.30.555526

**Authors:** Kylene P. Daily, Asmaa Badr, Mostafa Eltobgy, Shady Estfanous, Owen Whitham, Michelle H. Tan, Cierra Carafice, Kathrin Krause, Andrew McNamara, Kaitlin Hamilton, Samuel Houle, Spandan Gupta, Gauruv A. Gupta, Shruthi Madhu, Julie Fitzgerald, Abbey A. Saadey, Brooke Laster, Pearlly Yan, Amy Webb, Xiaoli Zhang, Maciej Pietrzak, Olga N. Kokiko-Cochran, Hazem E. Ghoneim, Amal O. Amer

## Abstract

Alzheimer’s Disease (AD) is the 6th leading cause of death in the US. It is established that neuroinflammation contributes to the synaptic loss, neuronal death, and symptomatic decline of AD patients. Accumulating evidence suggests a critical role for microglia, innate immune phagocytes of the brain. For instance, microglia release pro-inflammatory products such as IL-1β which is highly implicated in AD pathobiology. The mechanisms underlying the transition of microglia to proinflammatory promoters of AD remain largely unknown. To address this gap, we performed Reduced Representation Bisulfite Sequencing (RRBS) to profile global DNA methylation changes in human AD brains compared to no disease controls. We identified differential DNA methylation of CASPASE-4 (CASP4), which when expressed, can be involved in generation of IL-1β and is predominantly expressed in immune cells. DNA upstream of the CASP4 transcription start site was hypomethylated in human AD brains, which was correlated with increased expression of CASP4. Furthermore, microglia from a mouse model of AD (5xFAD) express increased levels of CASP4 compared to wild-type (WT) mice. To study the role of CASP4 in AD, we developed a novel mouse model of AD lacking the mouse ortholog of CASP4, CASP11, which is encoded by mouse *Caspase-4* (5xFAD/*Casp4^-/-^*). The expression of CASP11 was associated with increased accumulation of pathologic protein aggregate amyloid-β (Aβ) and increased microglial production of IL-1β in 5xFAD mice. Utilizing RNA sequencing, we determined that CASP11 promotes unique transcriptomic phenotypes in 5xFAD mouse brains, including alterations of neuroinflammatory and chemokine signaling pathways. Notably, *in vitro*, CASP11 promoted generation of IL-1β from macrophages in response to cytosolic Aβ through cleavage of downstream effector Gasdermin D (GSDMD). We describe a role for CASP11 and GSDMD in the generation of IL-1β in response to Aβ and the progression of pathologic inflammation in AD. Overall, our results demonstrate that overexpression of CASP4 due to differential methylation in AD microglia contributes to the progression of AD pathobiology, thus identifying CASP4 as a potential target for immunotherapies for the treatment of AD.

## Introduction

Alzheimer’s disease (AD) is a progressive neurodegenerative disorder and the 6^th^ leading cause of death in the United States (1). Currently, there are no reliable preventative or treatment methods for AD (2). Brain pathology in AD is characterized by extracellular senile plaques of amyloid-beta (Aβ) and intracellular neurofibrillary tangles of tau protein. An additional hallmark of AD is neuroinflammation, which contributes to the synaptic loss, neuronal death and symptomatic decline of AD patients (3).

Neuroinflammation in AD is coordinated by progressive changes in brain inflammatory cells, such as microglia and brain-associated macrophages (4). The changes in microglia responses are known to be regulated via epigenetic mechanisms (5,6). In particular, DNA methylation—an epigenetic mechanism that controls stable gene expression programs—is globally deregulated in neurodegenerative diseases such as AD (7,8). Therefore, it is plausible that the progressive inflammatory response seen in AD coincides with altered methylation status of critical immune effector molecules.

Amyloid beta (Aβ) is known to promote the production of inflammatory cytokines including IL-1β from microglia and macrophages (9), contributing to chronic long lasting sterile neuroinflammation in AD. For instance, the potent inflammatory cytokine IL-1β is implicated in AD pathogenesis by a variety of mechanisms. Multiple IL-1β genetic polymorphisms are associated with AD (10,11). IL-1β stimulates increased Aβ production from neurons and exacerbates neurofibrillary tangle formation (12–15). Thus, identifying the mediators involved in neuroinflammation and the production of IL-1β will provide mechanistic insight and potential diagnostic and therapeutic targets for AD.

Inflammatory mouse Caspase-11 (CASP11) and human orthologues Caspase-4 (CASP4) and Caspase-5 (CASP5) are the main drivers of noncanonical inflammasome activation which promotes release of IL-1β (16). We and others demonstrated that human CASP4 performs most of the functions of mouse CASP11, and for simplicity we will refer to them as CASP4 for human and CASP11 for mouse (17,18). In various biologic contexts, CASP4 promotes cleavage of Caspase-1 (CASP1) which activates IL-1β (16,19–21). Once cleaved, GSDMD forms pores in the plasma membrane, which results in the release of active IL-1β. The extensive formation of GSDMD pores can trigger inflammatory cell death and pyroptosis (22,23). GSDMD can be cleaved by active CASP1 or CASP11 according to the insult (20,21). Yet, it is currently unclear if CASP11 and GSDMD are required for CASP1 activation and inflammatory responses to Aβ. Importantly, CASP4 is upregulated in the brain of AD patients and correlates with disease progression and expression of risk genes (24,25). However, it remains largely unknown whether the increased expression of CASP4 is influenced by epigenetic factors or is associated with occurrence of hallmarks of AD pathobiology.

In this study, we profiled the global DNA methylation programs in brain tissues from AD patients and no disease controls. Importantly, we identified a unique DNA demethylation program upstream of the *CASP4* transcription start site in AD patients, that is correlated with an increased expression of CASP4 in human AD. To further understand the role of CASP4 in the development of AD, we developed a mouse model of AD (5xFAD, 5 familial Alzheimer’s disease mutations) lacking CASP11. We found that CASP11 drives Aβ deposition in male mice. Additionally, expression of CASP11 promoted release of IL-1β from microglia of 5xFAD mouse brains. Furthermore, transcriptomic profiling of the hippocampal tissue from 5xFAD/*Casp4^-/-^* mice revealed that CASP4 promotes neuroinflammation and chemokine signaling in 5xFAD mice. Finally, we performed *in vitro* analysis of inflammasome activation in response to Aβ introduced to cells by a cytosolic-delivery reagent in bone marrow-derived macrophages. We found that fibrillar Aβ activates the inflammasome response and that both CASP11 and NLR Family Pyrin Domain-Containing 3 (NLRP3) drive IL-1β activation and release. Our work positions CASP4 as a novel regulator of microglia inflammation in AD and demonstrates a molecular mechanism underlying its increased expression in AD brains.

## Results

### CASP4 upregulation is coupled with DNA demethylation events in human brains with Alzheimer’s disease

To better understand the DNA methylation deregulation during AD, we performed Reduced Representation Bisulfite Sequencing (RRBS) to profile global DNA methylation changes within frozen brain tissues at the temporal lobe (Brodmann area 38) from human AD brains and age- and sex-matched no disease (ND) controls. The temporal lobe is often the location where the characteristic spread of AD pathology including Aβ plaques starts (26). We found >4000 differentially methylated regions (DMRs) that were either hypomethylated or hypermethylated in AD versus ND brains (Fig 1A; Supplementary Data 1). Data shared through Gene Expression Omnibus with accession number GSE227194. To gain insights into the functional significance of these DMRs, we performed gene ontology (GO) enrichment analysis of the differentially methylated genes and found that hypomethylated DMRs in human brains with AD are enriched in biological processes regulating AD pathogenesis, such as Aβ formation (Fig 1B). In contrast, the hypermethylated DMRs in AD brains are enriched in biological processes regulating glutamate receptor signaling and neuronal cell-cell adhesion (Fig 1C). When we examined DNA methylation changes within *CASP4* loci, we identified a unique hypomethylated DMR located ∼ 350 bp upstream of the transcription start site at the CASP4 locus in AD patients as compared to patients without disease (p-value = 1.47E-6; n=5). To further validate the differential methylation state in AD brain samples, we designed a targeted epigenetic assay to assess DNA methylation levels at individual CpG sites in that genomic region (27). We found significant reduction in the average CpG methylation levels within this DMR in AD patients compared to non-dementia patients (Fig 1D). This DNA demethylation program was coupled with increased transcript and protein expression levels of CASP4 in human AD samples compared to non-dementia controls (Fig 1E-G). Notably, we found no difference in expression based on sex, when comparing the expression levels of CASP4 protein among human male and female brain samples (Fig 1F). These data indicate that CASP4 is upregulated in human brains with AD, which is epigenetically regulated, at least in part, by DNA demethylation programming.

**Figure 1.**
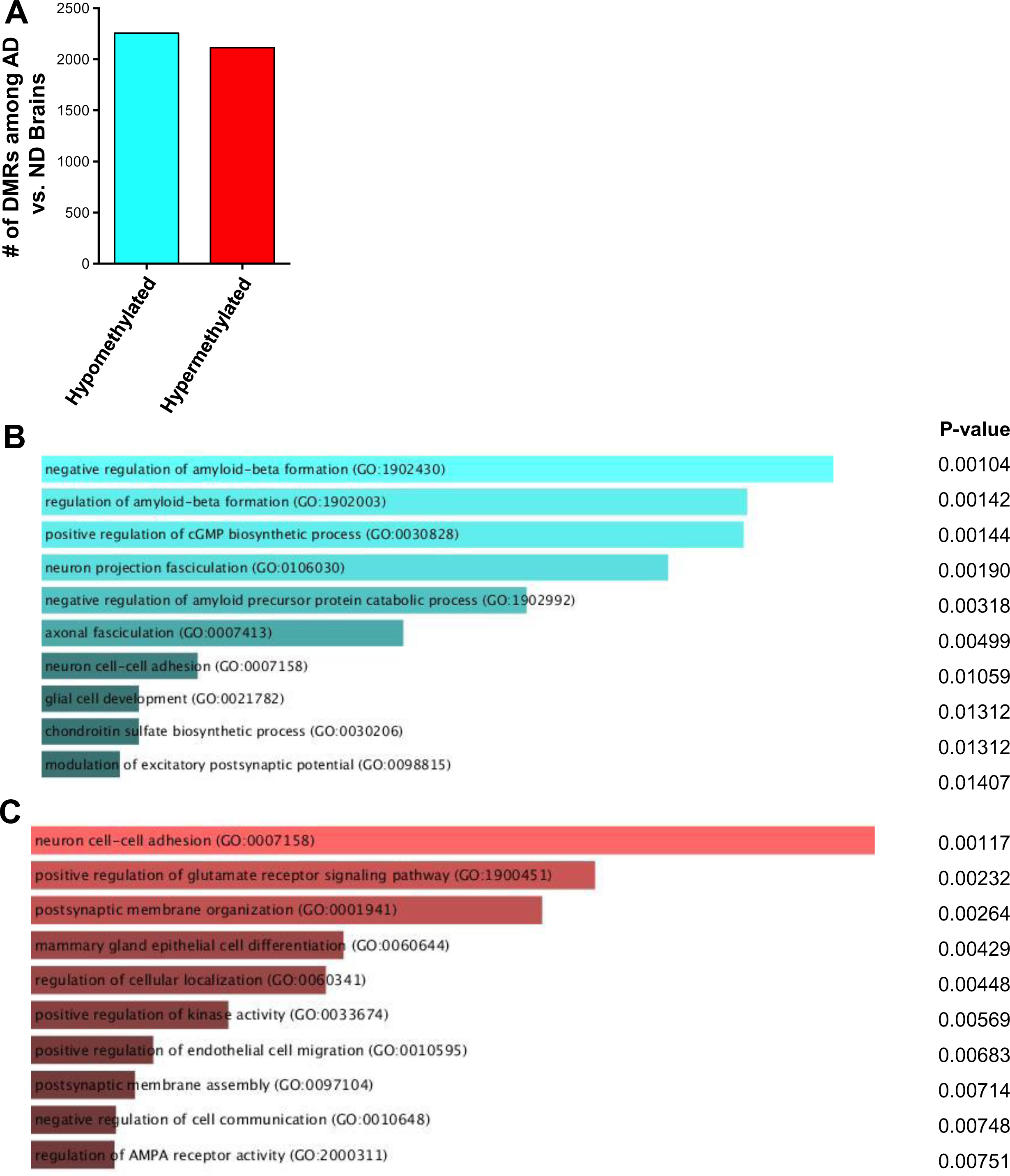
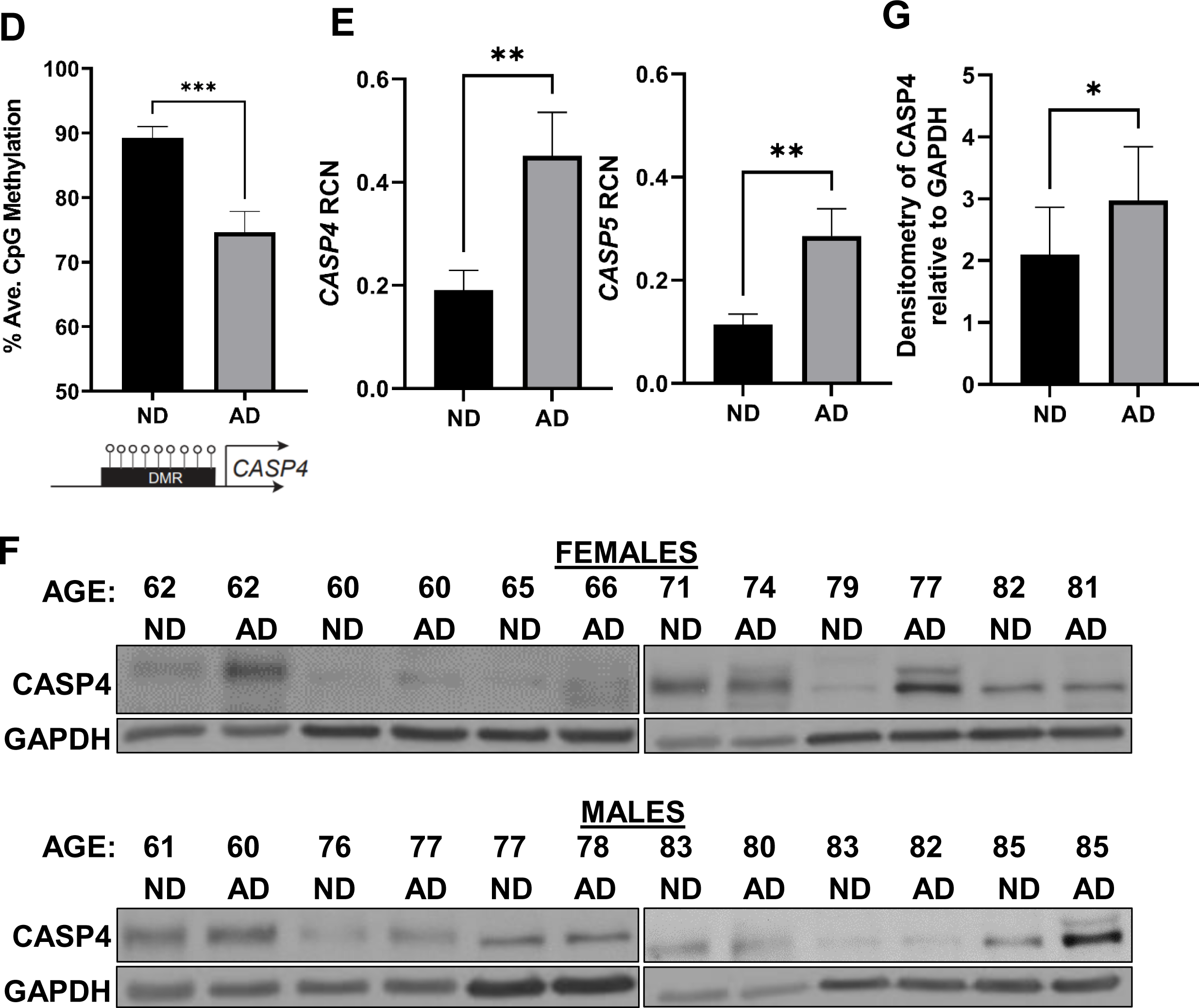
Human Alzheimer’s disease brains undergo distinct changes in DNA methylation including at the CASP4 locus. A) Numbers of differentially methylated regions (DMRs) identified by RRBS when comparing Alzheimer’s disease (AD) brains to non-dementia (ND) brains. B-C) Top GO biologic processes enriched for genes with hypomethylated (B) or hypermethylated (C) DMRs. D) Average CpG Methylation in temporal lobes of human Alzheimer’s disease (AD) and non-dementia (ND) controls for CASP4 differentially methylated region (DMR). Selected DMR is located ∼ 350 basepair upstream of the CASP4 transcription start site. Unpaired t-test with Welch’s correction (N=3 individual brain samples). *P ≤ 0.05, **P ≤ 0.01, ***P ≤ 0.001. E) Relative Copy Number (RCN) of CASP4 and CASP5 in temporal lobes of human AD and non-dementia controls determined by RT-PCR. Unpaired t-test (N=8). F) Immunoblot of CASP4 in temporal lobes of human AD and no disease (ND) controls. G) Densitometry of CASP4 relative to GAPDH above background for immunoblots in (F). Unpaired t-test (N=12).

### CASP11 is upregulated in 5xFAD mice specifically in microglia

We then determined if CASP11 expression was increased in the 5xFAD mouse model. The 5xFAD mouse model expresses 5 mutations detected in familial forms of AD under the Thy1 neuronal promoter (28). This model exhibits robust deposition of Aβ, starting at 2-months of age, followed by microgliosis and loss of cognitive function around 6 months (28). Expression of mouse *Casp4* which encodes CASP11 in the hippocampus of 7-month-old 5xFAD mice was analyzed by quantitative real-time PCR (RT-qPCR). 5xFAD mice express significantly more *Casp4* than age- and sex-matched wildtype (WT) littermates (Fig 2A). Importantly, there was no significant difference in the level of *Casp4* expression in 5xFAD male mice compared to female mice (Fig 2A). In other organs, *Casp4* is predominantly expressed in macrophages in response to DAMPs or PAMPs (29,30). To understand the cellular expression pattern of *Casp4* in the brain, we utilized a recently published pool of scRNA-seq datasets of immune cells in the mouse brain under homeostasis and analyzed the expression of *Casp4* expression levels in this scRNA-seq atlas (31). In agreement with our findings, we found that *Casp4* is mainly/highly expressed by microglia which also highly express CX3CR1 (Supplementary Figure 1). Since microglia are considered the main phagocytic immune cells of the brain, the expression of CASP11 in both 5xFAD microglia and non-microglia fractions was analyzed. Microglia were isolated from homogenized whole mouse brains by magnetic bead positive selection for CD11b to analyze expression of CASP11 by immunoblot. The non-microglia (CD11b-) fraction was also collected and contains all cell types other than microglia, such as neurons, endothelial cells, astrocytes, oligodendrocytes and ependymal cells as described previously (32). Microglia from 5xFAD mice expressed higher levels of CASP11 than age- and sex-matched WT mice (Fig 2B-C). There was no measurable expression of CASP11 in the combined non-microglia cells (Fig 2B). Therefore, we concluded that the expression of CASP11 is increased specifically in microglia of 5xFAD mice.

**Figure 2.**
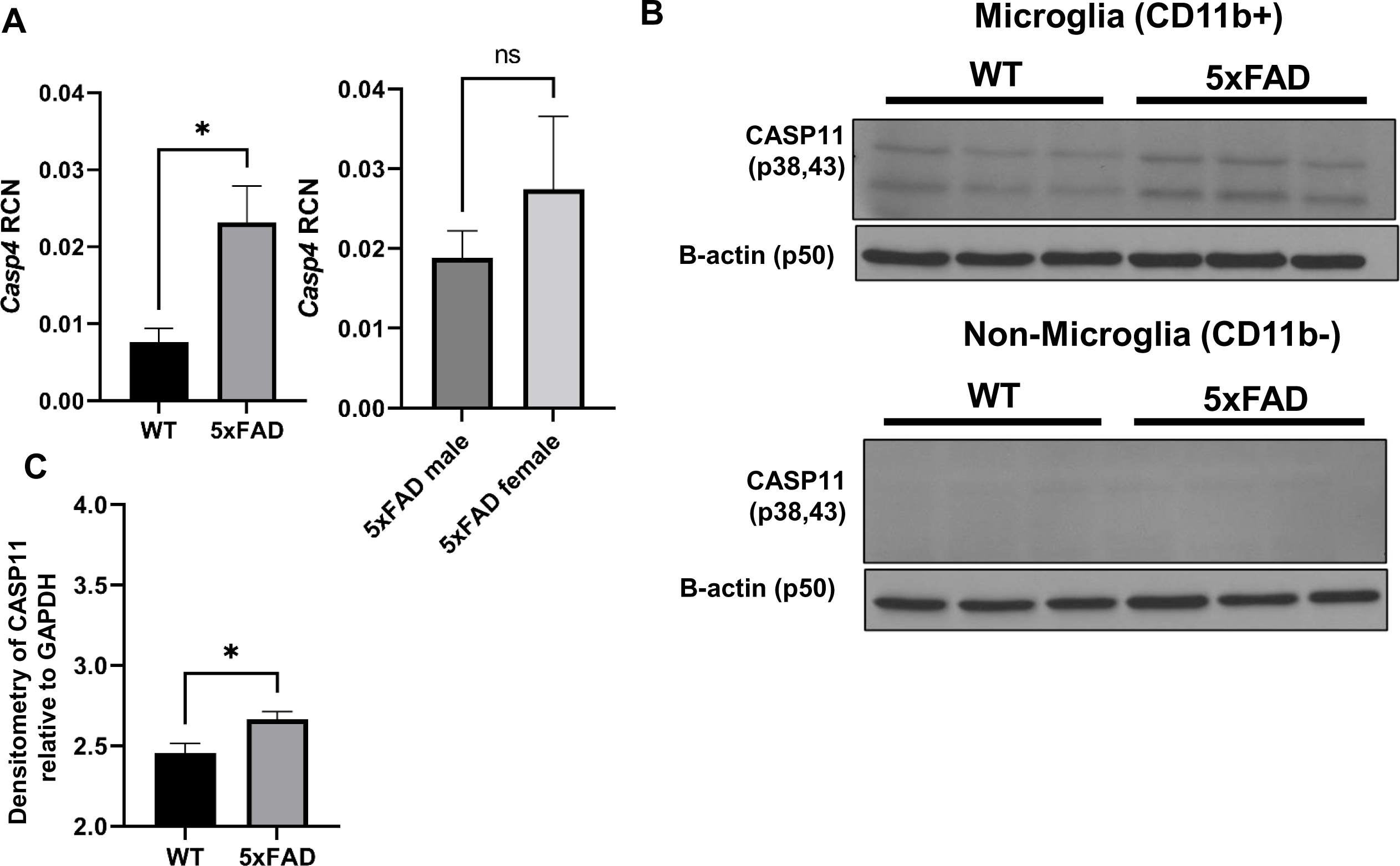
CASP11 exhibits increased expression in the 5xFAD microglia. A) Relative Copy Number (RCN) of *Casp4* in homogenized mouse brains from 7-month-old 5xFAD and age- and sex-matched wild-type (WT) littermate controls, and 5xFAD male mice compared to female mice determined by RT-PCR. RCNs are relative to housekeeping gene *GAPDH*, and multiplied by a factor of 100. Statistical analysis was completed by students t-test. B) Immunoblots for CASP11 and β-actin loading control in microglia (CD11b+) and non-microglia (CD11b-) fractions from 5-month-old WT and 5xFAD mice. C) Densitometry of CASP11 relative to GAPDH above background in microglia (CD11b+) for immunoblot shown in (B). (N=3) Statistical analysis completed by student’s t-test *P ≤ 0.05

### The expression of CASP11 drives brain pathology and inflammation in 5xFAD mouse brains

Since disease progression in the 5xFAD mouse correlates with increasing deposition of Aβ plaques (33), we characterized Aβ burden in the 5xFAD mice lacking *Casp4* gene (which encodes mouse CASP11 protein). To evaluate the role of CASP11 in 5xFAD brain pathophysiology, we crossed *Casp4^-/-^* mice with 5xFAD mice. We also maintained generation-matched WT/*Casp4^+/+^* (WT), WT/*Casp4^-/-^* (*Casp4^-/-^*) and 5xFAD/*Casp4^+/+^*(5xFAD) littermate mice. We quantified Aβ within the entire hippocampus of both 7-month-old male and female mice. To do so, the hippocampus was dissected, snap frozen and cryopulverized. We found Aβ is significantly reduced in male 5xFAD/*Casp4^-/-^* mice compared to 5xFAD (Fig 3A). However, in female mice we did not see a consistent reduction in Aβ by immunoblot (Fig 3B). As expected, we did not detect Aβ within the WT or *Casp4^-/-^*mice.

**Figure 3.**
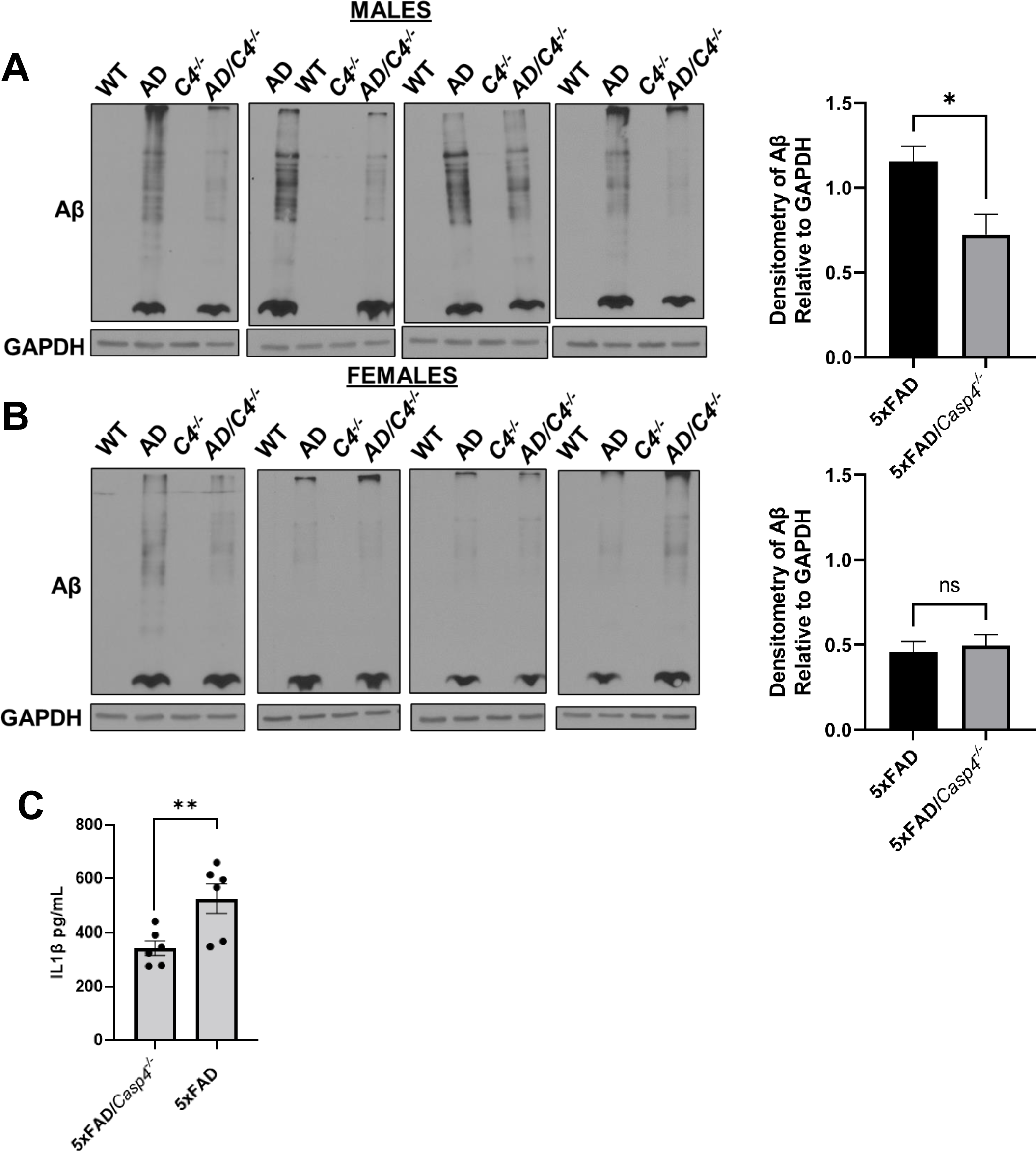
CASP11 deficiency leads to decreased Aβ deposition and reduced microglial-release of IL-1β in 5xFAD mice. Immunoblots for Aβ and GAPDH loading control in hippocampal lysate from 7-month-old littermate WT, 5xFAD, *Casp4^-/-^*and 5xFAD/*Casp4^-/-^* for A) male and B) female mice. Densitometry analysis of immunoblots for Aβ relative to GAPDH above background is displayed. Statistical analysis completed by student’s t-test *P ≤ 0.05. C) IL1β in cell culture supernatants from 5xFAD and 5xFAD/*Casp4^-/-^* microglia from 8-10-month-old male and female mice. Statistical analysis completed mixed effect analysis with Sidak multiple comparison test. (N=6) *P<0.05, **P<0.005.

It is well-described that AD is an inflammatory disease and that potent inflammatory cytokine IL-1β stimulates increased Aβ production (12–15). Therefore, we sought to determine if IL-1β release from microglia was exacerbated by the expression of CASP4. To do this, we isolated and cultured microglia from 8-10-month-old 5xFAD and 5xFAD/*Casp4^-/-^*mice for 24 hours after priming the inflammatory response with lipopolysaccharide (LPS) for 3 hours. We found that microglia from both male and female mice released more IL-1β when CASP4 was expressed (Fig 3C). Taken together, we concluded that the expression of CASP11 drives Aβ deposition in male 5xFAD mice, microglial release of IL-1β.

### CASP11 promotes neuroinflammation and chemokine production in 5xFAD mice

To define transcriptional underpinnings of the improved AD pathology in 5xFAD mice lacking CASP11, we performed bulk RNA-sequencing on the hippocampal region of 7-month-old age- and sex-matched 5xFAD and 5xFAD/*Casp4^-/-^* mice. We identified ∼860 differentially expressed genes among 5xFAD/*Casp4^-/-^ versus* 5xFAD brains (Figure 5A, Supplementary data file 2). Data shared through Gene Expression Omnibus with accession number GSE227193. The upregulated genes in 5xFAD/*Casp4^-/-^* brains were significantly enriched in various mitochondrial and metabolism related pathways, such as electron transport chain and oxidative phosphorylation (Fig 4B-C). In contrast, downregulated genes in the 5xFAD/*Casp4^-/-^* brains compared to 5xFAD brains were enriched in cell-cell adhesion and autophagy molecules (Fig 4D). Given the role of CASP11 in regulating inflammation and chemokine signaling, we further explored transcriptional changes in genes related to these pathways and indeed found multiple differentially expressed genes among the hippocampi of 5xFAD and 5xFAD/*Casp4^-/-^* mice within the neuroinflammation signaling, IL-8 signaling, and chemokine signaling pathways (Fig 4E-G). This led us to explore a role for CASP11 in inflammasome activation and IL-1β release *in vitro*.

**Figure 4.**
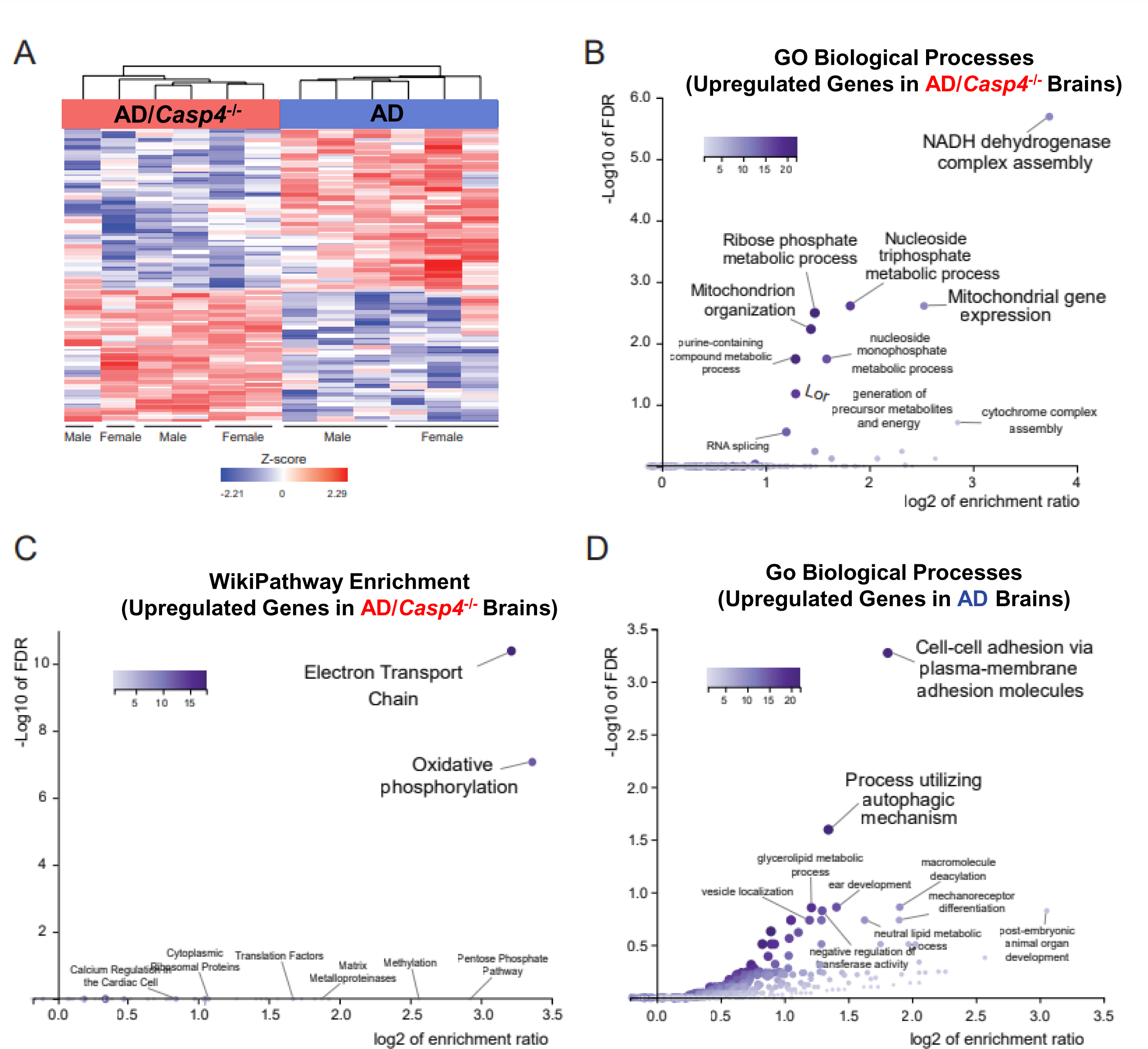
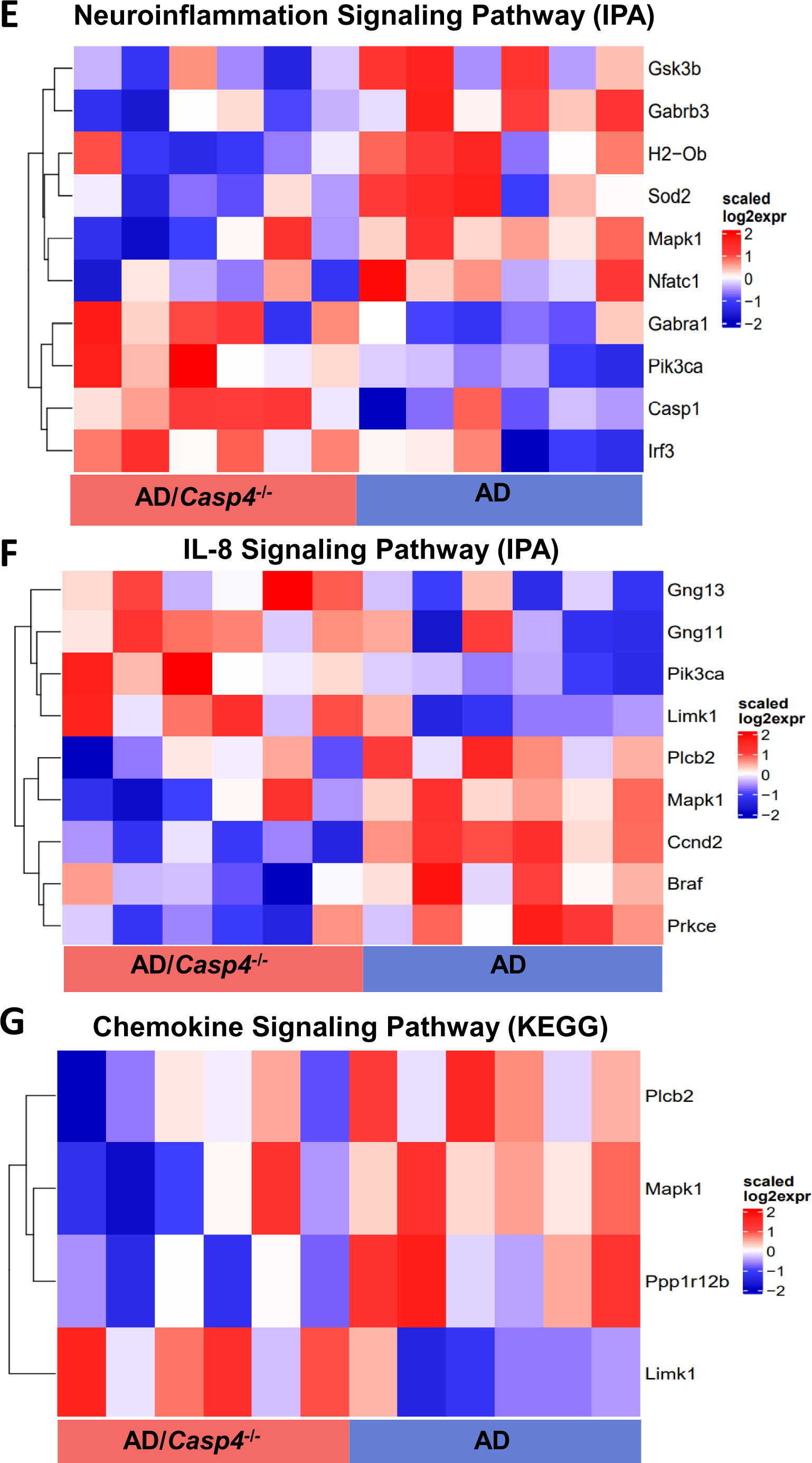
CASP11 promotes unique genetic phenotype including alterations of neuroinflammatory signaling pathways in the 5xFAD hippocampus. A) Heatmap showing differentially expressed genes, based on relative Z-score, in the hippocampus of 7-month-old 5xFAD (AD) and 5xFAD/*Casp4^-/-^*animals. B) GO-biological process enrichment analysis and (C) WikiPathway enrichment analysis for the upregulated genes in 5xFAD *Casp4^-/-^ versus* 5xFAD animals. D) GO-biological process enrichment analysis for genes upregulated in 5xFAD *versus Casp4^-/-^* 5xFAD animals. Heatmaps demonstrating differentially expressed genes: (E) neuroinflammation signaling pathway from IPA, (F) IL-8 signaling pathway from IPA, and (G) chemokine signaling pathway from KEGG.

**Figure 5.**
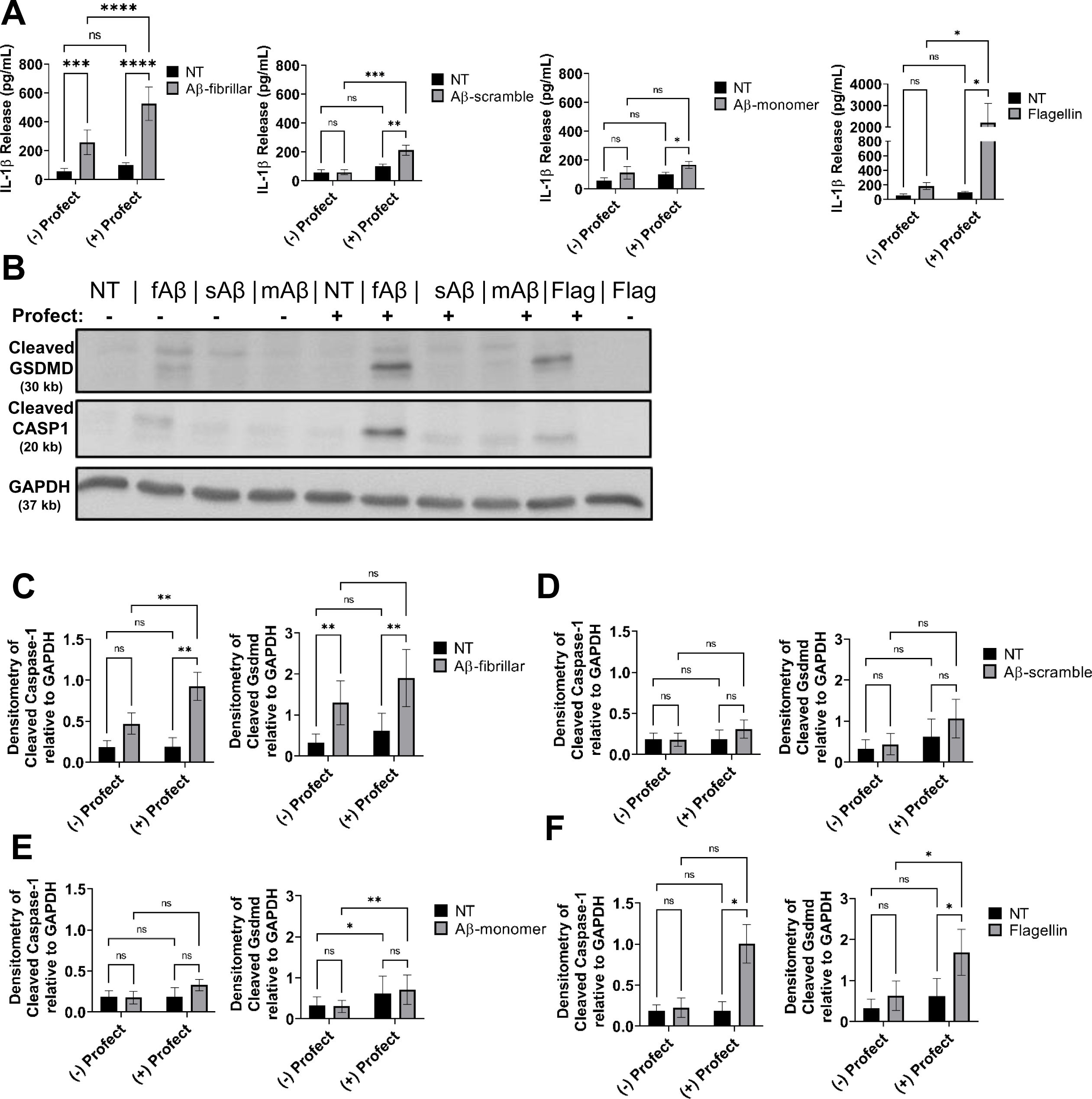
Profect-conjugated-fibrillar Aβ(1-42) stimulates release of IL-1β and cleavage of Gasdermin-D and Caspase-1. A) IL-1β release from LPS-primed mouse macrophages treated for 3 hours with 10µM fibrillar-Aβ (fAβ), 10µM scramble-Aβ (sAβ), 10µM monomer-Aβ (mAβ), 250ng/mL flagellin control or not treated (NT) with and without conjugation to cytosolic delivery reagent Profect (N=10). B) Representative immunoblots for cleaved GSDMD (30 kb), cleaved CASP1 (20 kb), and GAPDH (37 kb) from macrophages lysates treated as in (A). C-F) Densitometry analysis of immunoblots for cleaved GSDMD and cleaved CASP1 relative to GAPDH above background levels from LPS-primed mouse macrophages treated for 3 hours with C) 10µM fibrillar-Aβ with and without Profect, D) 10µM scramble-Aβ with and without Profect, E) 10µM monomer-Aβ with and without Profect, and F) 250ng/mL flagellin with and without Profect (N=6). Statistical analysis for (A) and (C-F) completed by 2way ANOVA Tukey’s multiple comparisons test with matching. For simplicity, graphs do not display p-values for all comparisons. *P ≤ 0.05, **P ≤ 0.01, ***P ≤ 0.001

**Figure 6.**
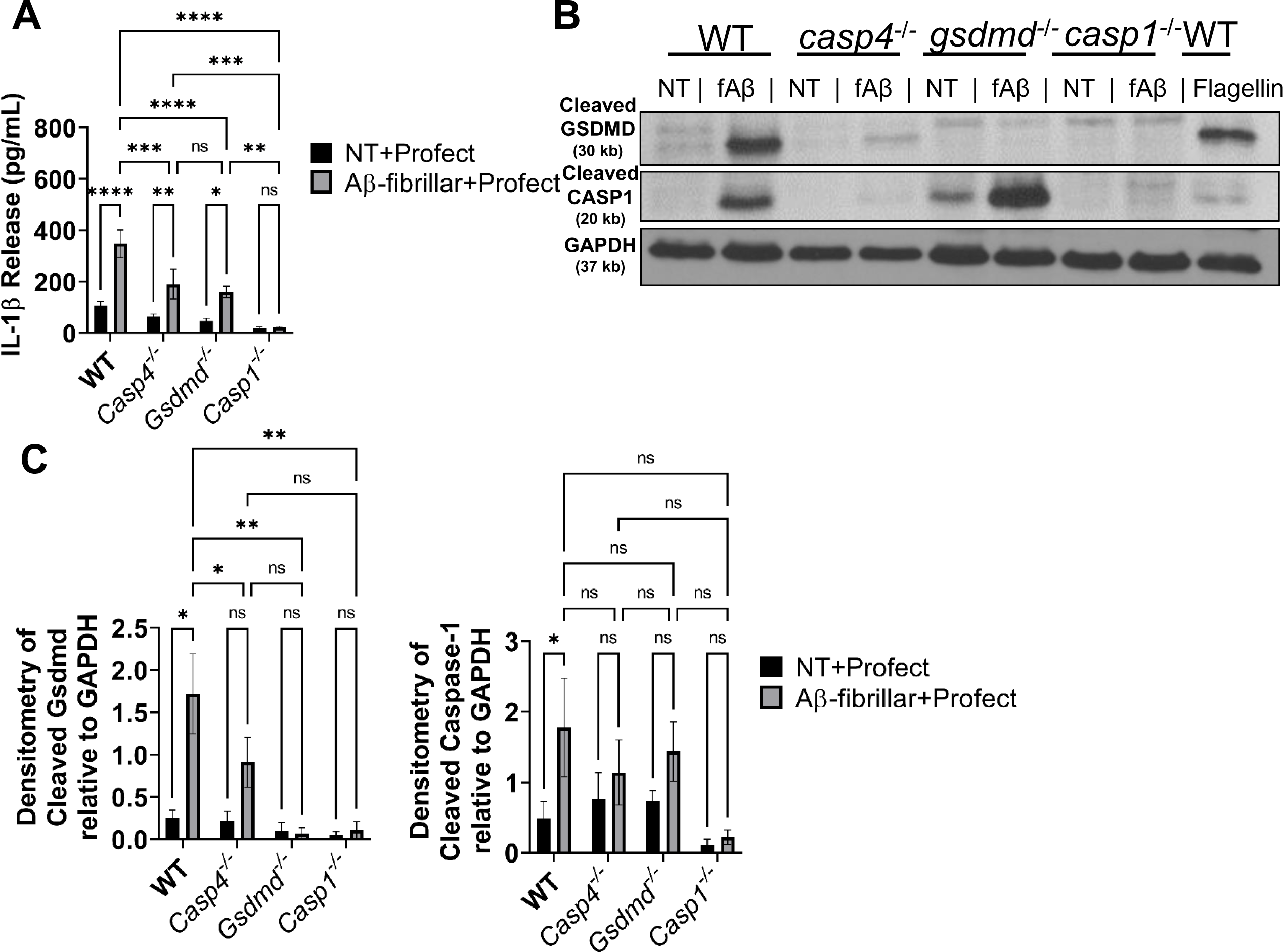
CASP11 promotes IL-1β release in response to Aβ by facilitating cleavage of GSDMD and CASP1. A) IL-1β release from LPS-primed macrophages from wild-type (WT), *casp4^-/-^*, *gsdmd^-/-^*, and *casp1^-/-^* mice treated for 3 hours with 10µM Profect-conjugated fibrillar-Aβ or Profect alone (NT+Profect) (N=9). B) Representative immunoblots for cleaved GSDMD (30 kb), cleaved CASP1 (20 kb), and GAPDH (37 kb) from macrophages lysates treated with Profect-conjugated fibrillar-Aβ (fAβ) or Profect alone (NT) as in (A). C) Densitometry analysis of immunoblots for cleaved GSDMD and cleaved CASP1 relative to GAPDH above background levels for macrophages treated as in (A) and (B) (N=8 for WT and *casp4^-/-^*, N=3 for *gsdmd^-/-^* and *casp1^-/-^*). Statistical analysis for (A) and (C) completed by 2way ANOVA Tukey’s multiple comparisons test. For simplicity, graphs do not display p-values for all comparisons. *P ≤ 0.05, **P ≤ 0.01, ***P ≤ 0.001, **** P ≤ 0.0001

As our male 5xFAD mice lacking CASP11 had dampened Aβ deposition, we also analyzed the data based on sex. For each group, only few genes that were differentially expressed between males and females were identified (Supplementary Table 1). The majority of these genes have previously been described as sexually dimorphic in the mouse hippocampus (*Xist, Eif2s3y, Uty, Ddx3y, Kdm5d*) (34). There was one gene, *Ptgds* encoding Prostaglandin D synthase, is reduced in males compared to females in the 5xFAD mice. Differential expression of *Ptgds* between males and females was not apparent in 5xFAD/*Casp4^-/-^* mice. Overall, we concluded that the involvement of CASP11 in driving neuroinflammation and Aβ production is influenced by sex, however more analysis for the exact mechanism is required.

### Cytosolic delivery of fibrillar-A**β** promotes maturation and release of IL-1**β** from immune cells without cell death

Activation of the inflammasome occurs within the cytosol where all components of the inflammasome machinery can interact upon activation (35). Macrophages and microglia are phagocytic cells that can internalize and traffic Aβ to lysosomes, although microglia in the AD brain exhibit defects in lysosomal function (36). The phagocytosis of increasing amounts of Aβ causes lysosomal rupture which results in release of the contents of the lysosome as well as cathepsin B into the cytosol (37,38). In this setting, cathepsin B serves as a signal for NLRP3 inflammasome activation. However, it remains unclear if the released Aβ in the cytosol could also activate the inflammasome as previous work did not differentiate the location of Aβ. Additionally, the contribution of CASP11 to IL-1β release in response to Aβ was not previously tested and we saw that CASP11 promoted microglial release of IL-1β. Therefore, we utilized a system in which Aβ is primarily delivered to the cytosol of mouse macrophages. We conjugated various forms of Aβ(1–42) to Profect (Targeting Systems), a reagent which transports proteins directly across the cell membrane into the cytosol of eukaryotic cells. We pre-treated wild-type (WT) macrophages with LPS to prime pro-IL-1β gene expression (39–41). Since soluble (monomeric), oligomeric and fibrillar forms of Aβ contribute to AD pathology (42), primed macrophages were treated with fibrillar-Aβ, an amino acid scrambled-Aβ control, and monomer-Aβ all with or without conjugation to Profect. The conjugation to Profect significantly increased IL-1β release in response to fibrillar Aβ. Scrambled-Aβ and monomer-Aβ did not induce IL-1β release without Profect conjugation (Fig 5A). We observed minimal, although significant, release of IL-1β in response to Profect-conjugated-monomeric or Profect-conjugated-scrambled-Aβ (Fig 5A). To confirm that Profect successfully transported proteins into the cytosol, we used Profect-conjugated-flagellin as a control (43). Profect-conjugated flagellin stimulated significant release of IL-1β (Fig 5A). These results indicate that fibrillar-Aβ promotes the release of IL-1β, and this effect is further enhanced when Aβ accesses the cytosol.

Previous work indicates that Aβ may prime the inflammasome through activation of toll-like receptors (TLRs) (41). Therefore, we determined if fibrillar-Aβ(1–42) treatment would prime the inflammasome response in macrophages. Macrophages were treated with fibrillar-Aβ with or without conjugation to Profect. LPS was used as a positive control. We utilized ATP as the second-step activator (20). LPS with ATP caused release of IL-1β, however ATP treatment of macrophages primed with fibrillar-Aβ with or without conjugation to Profect did not release IL-1β (Supplementary Figure 2A). This result indicates that fibrillar-Aβ(1–42) does not prime the inflammasome response in macrophages.

To show specifically that cytosolic fibrillar-Aβ activates the inflammasome response, we analyzed cell lysates for cleavage of CASP1 which is cleaved and activated by the inflammasome complex (44). We found that CASP1 is cleaved in response to fibrillar-Aβ conjugated to Profect and flagellin conjugated to Profect, but not in response to any of the other forms of Aβ (Fig 5B-F). Importantly, monomeric- and scrambled-Aβ conjugated to Profect did not cause CASP1 cleavage. Overall, these results indicate that cytosolic fibrillar-Aβ activates an inflammasome response leading to cleavage and activation of CASP1 and release of IL-1β.

The activation of CASP11 and/or CASP1 is accompanied by the cleavage of GSDMD and the release of the pore forming fragment (45). Cleaved GSDMD forms pores within the cellular plasma membrane allowing for release of mature IL-1β (20,46). To further determine the mechanism by which fibrillar-Aβ promotes IL-1β release, we analyzed cell lysates of WT macrophages and found that GSDMD is cleaved in response to fibrillar-Aβ, fibrillar-Aβ conjugated to Profect and flagellin conjugated to Profect, but not in response to any of the other forms of Aβ with or without Profect (Fig 5B-F). The excessive formation of GSDMD pores can lead to pyroptotic cell death leading to the release of IL-1β and lactate dehydrogenase (LDH) enzyme in some contexts (23,47,48). To determine whether pyroptotic cell death was occurring in response to fibrillar-Aβ, the release of LDH into the culture supernatants was analyzed. There was no difference in LDH release between no treatment (NT) and fibrillar-Aβ with and without Profect conjugation. Therefore, the cleavage of GSDMD and formation of pores in response to Aβ is not accompanied by cell death in response to fibrillar-Aβ.

### GSDMD is required for CASP11 and CASP1-mediated release of IL-1β following stimulation with fibrillar-Aβ

Earlier work suggests that the NLRP3 inflammasome mediates IL-1β release in response to Aβ, though the contribution of CASP11 to this process remains undetermined (37). We first evaluated if Profect conjugated-Aβ stimulates IL-1β release in an NLRP3 dependent manner in macrophages. We primed wild-type (WT) and *nlrp3^-/-^* macrophages with LPS and then treated with fibrillar-Aβ with or without conjugation to Profect. In response to fibrillar-Aβ conjugated to Profect, significantly reduced release of IL-1β in *nlrp3^-/-^*macrophages was observed (Supplementary Figure 3). These results indicate that cytosolic fibrillar-Aβ promotes the release of IL-1β via an NLRP3-dependent mechanism.

In addition to the canonical NLRP3 inflammasome, the noncanonical inflammasome can also activate CASP1 through CASP11. CASP11 cleaves GSDMD in some settings allowing for the release of IL-1β in CASP1 dependent and independent manners (20). However, the role of CASP11 in the inflammasome response to Aβ has not yet been differentiated. To determine if CASP11 mediates inflammasome activation in response to Aβ, we primed WT, *Casp4^-/-^, Gsdmd^-/-^*and *Casp1^-^* macrophages with LPS and then treated with fibrillar-Aβ with and without conjugation to Profect. In response to fibrillar-Aβ conjugated to Profect, significantly reduced levels of IL-1β were released from *Casp4^-/-^* and *Gsdmd^-/-^* and no IL-1β release from *Casp1^-/-^* macrophages when compared to WT (Fig 7A). These results indicate that CASP11 and GSDMD cleavage plays a role in promoting IL-1β release in response to fibrillar-Aβ in a CASP1-dependent manner.

To evaluate how CASP11 promotes IL-1β release, we analyzed the cleavage of CASP1 and GSDMD in cell lysates from LPS-primed WT, *Casp4^-/-^, Gsdmd^-/-^* and *Casp1^-/-^* macrophages treated with fibrillar-Aβ conjugated to Profect. CASP1 cleavage is significantly reduced in response to fibrillar-Aβ in *Casp4^-/-^* macrophages (Fig 7B-C), indicating that CASP11 expression promotes cleavage of CASP1. GSDMD can be cleaved by active CASP1 and/or CASP11 according to the insult (20). To determine if CASP11 and/or CASP1 are upstream of GSDMD cleavage, we analyzed cleavage of GSDMD in the absence and presence of Casp4/11. We found that GSDMD cleavage was decreased in both *Casp4^-/-^*and *Casp1^-/-^* macrophages in comparison to WT (Fig 7B-C), indicating that both CASP11 and CASP1 promote GSDMD cleavage in response to fibrillar Aβ. Overall, these results indicate that both the noncanonical inflammasome and NLRP3 inflammasome mediate IL-1β release following exposure to Aβ via GSDMD.

## Discussion

In this study we demonstrated that CASP4 (CASP11) is a driver of chronic neuroinflammation in AD in response to Aβ. Our work identified that CASP4 (CASP11) is upregulated in the temporal lobe (Brodmann area 38) of AD patients compared to age- and sex-matched non-dementia controls. According to a previous study which analyzed expression data from over a thousand AD patient samples, *CASP4* gene expression is also increased in the cerebellum, dorsolateral prefrontal cortex and visual cortex of AD patients (24). Notably, *CASP4* expression positively correlates with increased expression of *CASP1,* as well as AD risk factor genes *TREM2, CR1,* and *TYROBP* which are implicated in microglia mediated inflammation (24). An additional study found that *CASP4* expression is increased in the hippocampus of AD patients and correlates with clinical disease progression (25). Our work indicates that epigenetic mechanisms may underly the increased expression of *CASP4* in the AD brain.. We also discerned the contribution of members of the non-canonical inflammasomes to IL-1β release in response to Aβ.

In the AD field, several studies have completed analysis of the methylation status of specific candidate genes that are implicated in AD pathogenesis (49). These include *APOE*, *BDNF*, *TREM2*, and glycogen synthase kinase 3 beta (*GSK3*β). Genome wide DNA methylation studies on AD patients indicated that numerous pathways such as neurogenesis, amyloid generation, and inflammatory responses are implicated (49). It is currently unclear whether key genes implicated in AD pathogenesis are associated with hypermethylation, hypomethylation or a mix of both. According to our data, effectors involved in Aβ generation may experience global hypomethylation leading to increased production of Aβ which is in agreement with prior studies (49). We also found that other critical neuropathways, such as cell adhesion and glutamate signaling, undergo hypermethylation as reported previously (49). Although globally altering the methylation status was previously suggested as a therapeutic option in AD, these recent combined data suggest that benefits maybe counteracted by unfavorable effects.

The precise mechanism underlying differential methylation of *CASP4* in AD patients remains unknown and warrants further evaluation. DNA methylation, denoted as 5-methylcytosine (5mC), is a well-characterized and highly stable epigenetic mechanism (50–52). Differentiating mechanisms underlying demethylation programming has been crucial to understand this epigenetic mark of DNA hypomethylation (7). Potential pathways include loss of maintenance DNA methylation programs, termed passive demethylation, as well as nucleotide and base excision repair pathways, or active demethylation (7,8,53). DNA methyltransferase 1 (DNMT1) is responsible for maintenance of DNA methylation. Notably, DNMT1 as well as other methylation factors are decreased in Alzheimer’s disease brain tissue, which could lead to passive demethylation (8). DNA demethylation can also be initiated by active mechanisms include a mediator step of oxidation of 5mC into 5-hydroxymethylcytosine (5hmC) by the ten-eleven translocation (TET) proteins (7). Notably, 5hmC is especially enriched in the brain (54). Bilsufite sequencing as performed here converts both 5mC and 5hmC to uracil, thereby analyzing both as methylated sites (55). In general, 5hmC is associated with increased gene activity, although this may depend on other factors (56). A recent study found hundreds of differentially hydroxymethylated 5hmC regions associated with Aβ plaques and neurofibrillary tangles in human AD patients yet *CASP4* was not identified as a differentially hydroxymethylated region (57). According to our data, hypomethylation of DNA upstream of the *CASP4* transcription start site may stably enhance expression of CASP4 in the AD brain. Our data points to an epigenetic mechanism underlying the exacerbated IL-1β release and inflammasome activation seen in AD which improves our understanding of neuroinflammation in this disease. As CASP4 is primarily expressed in microglia, microglia-targeted analysis of DNA demethylation programs in AD would be beneficial to further understand the mechanism underlying increased *CASP4* expression. Accordingly, microglia-targeted methylation therapies can be effective and accompanied by less off target-mediated side-effects.

We demonstrated by multiple methods that CASP11 protein is primarily expressed in microglia and is increased in 5xFAD microglia. In support of this finding, according to the online resource from the Barres group, *Casp4* mRNA is predominantly expressed in macrophages and microglia, with a lower level of expression in endothelial cells in healthy mouse brains (29,30). Microglia exhibit various roles in AD progression including generation of IL-1β, which drives production and seeding of Aβ plaques, exacerbates neurofibrillary tangle formation and promotes tissue damage and synaptic dysfunction in AD (12–15,37,58,59). However, the mechanism underlying the elevated chronic production of IL-1β by microglia in AD and a potential role for CASP4 were still unclear.

We were able to further analyze the contribution of CASP4 to AD disease progression by generating a 5xFAD mouse lacking murine *Casp4* expression. We found that CASP11 drives neuroinflammation and Aβ deposition. The RNAseq analysis on the hippocampus from 5xFAD and 5xFAD/*Casp4^-/-^* mice revealed differences in genes encoding important AD pathogenesis and neuroinflammatory regulators. We observed significantly decreased expression of *GSK3B* in 5xFAD/*Casp4^-/-^*mice (Fig 4A). *GSK3B* encodes Glycogen synthase kinase 3β (GSK3β), which contributes to increased Aβ production, hyperphosphorylation of tau, and microglia activation (60). Consequently, inhibition of GSK3β in mice reverses AD pathology (60). We also observed a decrease in *MAPK1* which encodes Mitogen-Activated Protein Kinase 1 (MAPK1) in 5xFAD/*Casp4^-/-^* (Fig 4E-G). The MAPK family are crucial regulators of neuroinflammatory processes including generation of inflammatory cytokines (61). Inhibition of MAPK1 in a rat model of AD improved cognitive function (62). Our extensive work in CASP4 biology demonstrated consistently that CASP4 promotes inflammatory cell migration in various contexts (63,64). Prior work also demonstrated that expression of CASP4 promotes microglia clustering around Aβ plaques and increased inflammation in the brain of AD mice (24). It is therefore likely that CASP4 promotes increased immune cell migration and activation in AD brains.

Few reports have characterized IL-1β activation in response to Aβ (37,59,65), but our study evaluated the role of both CASP11 and GSDMD as new molecular players in the highly complex multi-protein process required for IL-1β activation and release in AD. Activation of the inflammasome, which promotes maturation and release of IL-1β, requires a priming signal and an activating signal (63). The activating signal is a cytosolic pathogen-associated molecular pattern (PAMP) or a danger-associated molecular pattern (DAMP) (41). CASP11 is known to potentiate inflammasome responses leading to GSDMD pore formation which drives the release of active cytokines (66–68). Several studies demonstrated that NLRP3 promotes pathology in a mouse model of AD (59). As such, NLRP3 is widely accepted as a potential therapeutic target, although drug-design against this complex structure has proven to be difficult (69,70). We demonstrate that CASP4 promotes cleavage of both CASP1 and GSDMD *in vitro* in macrophages treated with Profect-conjugated fibrillar Aβ. Currently, the established mechanism of Aβ-induced inflammasome activation is via an indirect mechanism that can be explained through several possibilities. It has been shown that Aβ promotes lysosomal damage leading to release of cathepsin B and subsequent activation of the NLRP3 inflammasome (37). Supporting this mechanism, here we demonstrated that cytosolic Aβ directly stimulates an inflammasome response. It is plausible that ROS production associated with Aβ phagocytosis also activates the NLRP3 inflammasome (37,38,65).

IL-1β is known to stimulate the production of *AMYLOID PRECURSOR PROTEIN* (*APP*) and enhance the activity of enzymes that generate Aβ (12,13). Therefore, CASP4 may promote Aβ plaque generation by enhancing IL-1β production and release. Additionally, the release of apoptosis-associated speck-like protein containing a CARD (ASC) as a by-product of inflammasome activation serves as a seeding point for the formation of Aβ plaques (71). Aβ bound to ASC is also more toxic to microglia and promotes increased microglia death and dysfunction (72). CASP4 increases the release of ASC through GSDMD pores which allows for more Aβ plaques to form thereby promoting continued activation of microglia. It is also possible that CASP4 alters the ability of microglia to clear Aβ via modulation of actin dynamics as observed in response to bacterial infection of macrophages (73–76). Reduced Aβ burden in 5xFAD mice lacking CASP4 may therefore be due to reduced inflammation which reduces the production of Aβ by neuronal cells.

CASP4 expression is likely regulated in part by methylation status. Modulation of epigenetic mechanisms has been suggested as a potential strategy for the treatment of AD, especially with the success of this approach in some cancers (77–79). Studies testing therapeutic DNA methylating and DNA demethylating agents for AD could utilize CASP4 expression as an epigenetic marker of microglia activity. In conclusion, we provide mechanistic evidence that CASP4 is a regulator of neuroinflammation in AD with potential as a therapeutic target.

## Materials and Methods

### Human Samples from NIH Biobank

Human temporal lobe brain tissue sections (Brodmann area 38) with AD and age-matched controls without dementia (non-dementia, ND) were obtained from the NIH biobank in accordance with our institution’s IRB. These snap-frozen tissue samples were used for all TRIzol homogenization, RT-qPCR and immunoblot analysis. Range of hours until freeze, age, race and sex are in Supplementary Table 2.

### Reduced Representation Bisulfite Sequencing and Analysis

Five samples from AD and age-matched non-dementia controls were prepared for RRBS analysis. DNA methylation data were generated for all samples using the reduced representation bisulfite sequencing (RRBS) method (80). Briefly, NEXTflex Bisulfite Library Prep Kit for Illumina Sequencing (Bioo Scientific, PerkinElmer, MA) was used to convert the study sample genomes to bisulfite-converted genomes. Illumina sequencing libraries were prepared using Zymo EZ-DNA methylation kit. Bisulfite converted libraries were sequenced to 35 – 40 million PE-150×150bp clusters per samples using HiSeq 4000 Sequencer (Illumina, CA).

Raw read QC assessed with FastQC v0.11.5 before trimming and after trimming. Trimming performed with TrimGalore v0.4.5 specifying RRBS and Illumina with a quality cutoff of 20. Quality and adapter trimmed reads aligned to bismark generated indexes for GRCm38 with bismark v0.22.1 using bowtie2 v2.3.5.1 with options “--N 1 --L 15 -D 50 --score_min L,-0.6,-0.6 -p 10 --X 600” Methylations extracted using bismark_methylation_extractor. Reports generated with bismark2report. Alignment bam sorted and indexed with samtools v1.9. Differential methylation of regions performed with R package DSS running DMLtest with group1 as Control and group2 as Alzheimers and callDML and callDMR with option p.threshold=0.05. Regions annotated with R package AnnotationDbi and human database org.Hs.eg.db DMR results filtered with a threshold of 50% change in methylation ratio and *P-value <0.01*, with >5 reads in at least 2 samples among human brain samples. Gene Ontology biological process enrichment analysis was performed using Enrichr for genes with hypomethylated or hypermethylated DMRs (81).

### Targeted DNA Methylation Analysis of the CASP4 Locus

Genomic DNA was extracted from snap-frozen brain samples using the PureLink® Genomic DNA Mini Kit (cat no: K182002, Invitrogen), and then bisulfite-treated using the EZ DNA Methylation-Direct Kit (Zymo) to convert all unmethylated cytosines to uracil. Locus-specific PCR was performed on the bisulfite-converted DNA using the primers specific for the DMR at the *CASP4* locus: *FP (ggaatttagtttttgatttggggg) and RP (cccacctaaaaaaacaatctaacc).* The amplicon DNA size was confirmed by gel electrophoresis and purified using the Zymoclean Gel DNA Recovery Kit (Zymo). DNA sequencing library of the purified CASP4 PCR amplicons was prepared using the Native Barcoding Kit (cat no: EXP-NBD104) and Ligation Sequencing Kit (cat no: SQK-LSK109) (Oxford Nanopore Technologies). Sequencing was performed on the prepared library using an R9 flow cell and MinION device (Oxford Nanopore Technologies). The resultant FASTQ files from the MinION sequencing were extracted for downstream analysis of CpG methylation at the amplified DMRs. In brief, the FASTQ files were quality controlled by first removing the multiplexing barcodes using Porechop v0.2.4 (github.com/rrwick/Porechop) with the default parameters and then filtering by read quality (-q 10) and length (-l 200) using NanoFilt v2.6.0. (82). The quality-controlled reads were then mapped against the PCR target amplicons (-x map-ont) using minimap2 v2.17-r941 (83). For each amplicon, the different types of bases mapped at each position in the bam files (count --bases -w 1) were counted using igvtools v2.8.0.(84) The CpG sites were automatically extracted from the PCR target sequences (locate -i -P -p cg) using SeqKit v0.12.0 (85) and custom bash scripts were used to extract the base distribution from igvtools at these specific CpG locations and tabulate them per amplicon.

### Mice

C57BL/6 wild-type (WT) mice and 5xFAD mice (stock #034840) were obtained from the Jackson Laboratory (Bar Harbor, ME, USA). C57BL/6 WT mice were obtained from the Jackson Laboratory (Bar Harbor, ME, USA). *Casp4^-/-^* mice were generously provided by Dr. Yuan at Harvard Medical School, Boston, MA, USA and are now available from the Jackson Laboratory (024698).(18) *Casp-1*^-/-/Casp11Tg^ mice were kindly provided by Dr. Vishva Dixit at Genentech, San Francisco, CA, USA. *Gsdmd^-/^*^-^ mice and *nlrp3^-/-^* bones were provided by Dr. Thirumala-Devi Kanneganti at St. Jude Children’s Research Hospital, Memphis, TN, USA. All mice were housed in a pathogen-free facility, and experiments were conducted with approval from the Animal Care and Use Committee at the Ohio State University (Columbus, OH, USA).

### Creation of 5xFAD/Casp4 strain

The 5xFAD (AD) (B6SJL-Tg (APPSwFlLon, PSEN1* M146L* L286V) 6799Vas /Mmjax) mouse is a double transgenic APP/PS1 mouse model that co-expresses five AD mutations leading to accelerated plaque formation and increased Aβ42 production. AD mouse model over-expresses APP with K670N/M671L (Swedish Mutation), I716V (Florida mutation), and V717I (London mutation), PS1 with M146L and L286V mutations. These mice accumulate high levels of intra-neuronal Aβ-42 around 1.5 months of age with amyloid deposition around 2 months (28,86). The 5xFAD mice were crossed with *Casp4^-/-^* mice for 5-6 generations for use in tissue or behavioral assessments. 5xFAD mice were maintained hemizygous by crossing littermate mice not carrying the transgene with those carrying one copy of the transgene in order to generate the full *Casp4* knockout. Genotyping was carried out for *Casp4* mutant and 5xFAD transgene as specified by the Jackson Laboratory.

### Preparation of mouse hippocampus

Mouse brains were dissected following euthanasia by CO_2_. The entire hippocampus was immediately dissected on ice as described previously (87) and snap frozen in liquid-nitrogen and stored at −80°C. The hippocampus was cryopulverized into a fine powder on a liquid-nitrogen frozen BioPulverizer (cat no: 59012MS, BioSpec). The powder was mixed well and collected in separate tubes for future analysis. For immunoblot and RT-qPCR analysis, brains were lysed in TRIzol. For RNAseq analysis, brains were lysed as with lysis buffer from PureLink™ RNA Mini Kit (Invitrogen, 012183025) by manufacturers instructions.

### Microglia isolation

5-month-old, sex-matched 5xFAD and WTAD mice were utilized for microglia isolation to analyze expression of CASP11 by immunoblot. Microglia were isolated by MACS neural dissociation kit (Miltenyi Biotec, 130-107-677) followed by CD11b magnetic bead (Miltenyi Biotec, 130-093-634) isolation technique to positively select for brain microglia expressing the pan-microglia marker CD11b as has been described before (32,88). Microglia (CD11b+) and non-microglia (CD11b-) fractions were collected. Cells were pelleted and washed one time with PBS prior to lysing with TRIzol.

### Cell culture

Primary bone marrow-derived macrophages were derived from mice as previously described (75). Briefly, tibias and femurs were flushed with IMDM media (Thermo Fisher Scientific, 12440053) supplemented with 10% heat inactivated fetal bovine serum (FBS, Thermo Fisher Scientific, 16000044), 50% L cell-conditioned media, 0.6× MEM Non-Essential Amino Acids (Thermo Fisher Scientific, 11140050), and 0.1% penicillin and streptomycin (Thermo Fisher Scientific, 15140122). Cells were cultivated at least 6 days at 37°C in a humidified atmosphere containing 5% CO2.

Microglia were isolated from the brains of 8-10 months old mice and then cultured in the same media listed for macrophages. For measurement of IL-1β in cell culture supernatant, microglia were primed with 100ng/mL LPS for 3 hours and then supernatant was collected after 24 hours of culture.

### A**β** peptides and fibrillization

Aβ peptides in powder form (Beta-Amyloid (1–42) cat no: AS-20276; Scrambled-beta-Amyloid (1–42) cat no: AS-25382, Anaspec) were initially resuspended in 1% NH_4_OH (cat no: AS-61322, Anaspec) for 15 min to dissolve any pre-formed aggregates per manufacturer instructions, and then diluted to a final concentration of 0.05% NH_4_OH in water prior to fibrillization. Aβ peptides were converted to the fibrillar form of Aβ by incubating monomeric human Aβ(1–42) at 220 μM in water at 37°C for 3 days prior to use as previously described (89). Scrambled Aβ was also treated for fibrillization, although this should not form aggregates.

### In vitro treatment of primary macrophages

Macrophages were cultivated in IMDM media supplemented with 10% FBS at 1×10^6^ cells per well in a 12-well plate. Macrophages were primed for 3 hours with 100 ng/mL Ultrapure LPS, E. coli 0111:B4 (Invivogen). 10 µM Aβ peptides were conjugated to Profect-P1-lipid based protein delivery reagent (cat no: 0041, Targeting Systems) in 100µL serum free RPMI for 20 minutes prior to gentle resuspension in a final volume of 425 µL. Negative control Profect was incubated without any Aβ peptide. Macrophages were washed 3 times with serum free RPMI to remove serum and free LPS prior to treatment with Profect-complexes. Macrophages were treated with 400 µL Profect complexes for 3 hours. Supernatant was collected for analysis by ELISA and macrophages were lysed in TRIzol reagent. For experiments utilizing ATP, macrophages were not primed with LPS and instead treated first with Profect-complexes for 3 hours or LPS as a positive control, and then with or without 5 mM ATP (cat no: A6419, Sigma Aldrich) for 30 minutes.

### Immunoblot

Protein extraction was performed using TRIzol reagent (Thermo Fisher Scientific) according to the manufacturer’s instructions. Briefly, after phase separation using chloroform, 100% ethanol was added to the interphase/phenol-chloroform layer to precipitate genomic DNA. Subsequently, the phenol-ethanol supernatant was mixed with isopropanol to isolate proteins. The Bradford method was used to determine protein concentrations. Equal amounts of protein were separated by 13.5% SDS-PAGE and transferred to a polyvinylidene fluoride (PVDF) membrane. Membranes were incubated overnight with antibodies against human CASP4 (cat no: M029-3, MBL), mouse Caspase-11 (cat no: NB120-10454, Novus Biologicals), human Aβ (D54D2) (cat no: 8243S, Cell Signaling Technology), mouse Gadermin-D (cat no: ab209845, Abcam), mouse Caspase-1 (cat no: AG-20B-0042-C100, AdipoGen), β-actin (cat no: 3700S, Cell Signaling Technology) or GAPDH (cat no: 2118, Cell Signaling Technology). Corresponding secondary antibodies conjugated with horseradish peroxidase (cat no: 7074 Rabbit, cat no: 7076 Mouse, cat no: 7077 Rat, Cell Signaling Technology) in combination with enhanced chemiluminescence reagent (cat no: RPN2209, Amersham) were used to visualize protein bands. Densitometry analyses were performed by normalizing target protein bands to their respective loading control using ImageJ software as previously described (73).

### Measurement of IL-1β

The level of IL-1β in macrophage culture supernatants was measured by R&D Systems DuoSet ELISA Development Systems (murine IL-1b, DY401) according to the manufacturer’s instructions. The level of IL-1β in microglia culture supernatants was measured by Procartaplex Mouse Cytokine and Chemokine kit (cat no: EPXR260-26088-901, Thermo Fisher Scientific) using the MAGPIX plate reader according to the manufacturer’s instructions.

### LDH Release

LDH release from macrophages treated with Aβ profect complexes was measured using the CytoTox-ONE Homogeneous Membrane Integrity Assay (cat no: G7891, Promega) according to the manufacturer’s instructions. Aβ-induced LDH release [%] = ((test sample)/(high control))*100.

### Quantitative real-time PCR (RT-qPCR)

Total RNA was isolated from cryopulverized tissue lysed in TRIzol (cat no: 15596026; Invitrogen Life Technologies). Chloroform (cat no: 268320010, Fisher Scientific), isopropanol (cat no: BP2618212, Fisher Scientific), and glycogen (cat no: 10814010, Fisher Scientific) were used to isolate total RNA and its concentration was measured by Nanodrop. The expression of genes was determined as previously described and expressed as relative copy numbers (RCN) (90). RCNs are relative to housekeeping genes GAPDH and CAP-1, and multiplied by a factor of 100. Ct values of each *gene* were subtracted from the average Ct of the internal control. The resulting ΔCt was used in the equation: RCN= (2^-ΔCt^)*100. The following primer sets were used: Human *CASP4* (FP-cacaacgtgtcctggagaga, RP-acttcctctaggtggcagca), human *CASP5* (FP-agtcagtgctgagggcattt, RP-ccctctaggatgccatgaga), mouse *Caspase-4* (FP-catcactagactcatttcctgctt, RP-ctggaatttcaggaatagaatgtg).

### RNA-sequencing and data analysis

Total RNA was extracted from cryopulverized hippocampal tissue by PureLink™ RNA Mini Kit (Invitrogen, 012183025) according to the manufacturer’s instructions following lysis with the provided lysis buffer. RNA cleaning and concentration was done using Zymo Research, RNA Clean & Concentrator™-5 kit (cat# R1015) following the manufacturer’s protocol. Fluorometric quantification of RNA and RNA integrity analysis was carried out using RNA Biochip and Qubit RNA Fluorescence Dye (Invitrogen). cDNA libraries were generated using NEBNext® Ultra™ II Directional (stranded) RNA Library Prep Kit for Illumina (NEB #E7760L). Ribosomal RNA was removed using NEBNext rRNA Depletion Kit (human, mouse, rat) (E #E6310X). Libraries were indexed using NEBNext Multiplex Oligos for Illumina Unique Dual Index Primer Pairs (NEB #644OS/L). Library prep generated cDNA was quantified and analyzed using Agilent DNA chip and Qubit DNA dye. Ribo-depleted total transcriptome libraries were sequenced on an Illumina NovaSeq SP flow cell (paired-end 100×100 bp format; 35-40 million clusters, equivalent to 70-80 million reads. Library preparation, QC, and sequencing was carried out at Nationwide Children’s Hospital genomic core.

RNA sequencing data processing and analysis were performed by the Biomedical Informatics Shared Resource (BISR) group at the Ohio State University using previously published pipeline (91). Briefly, raw reads were aligned to mouse reference genome GRCm38 with HISAT2 v2.1.0 (92). Gene wise counts were generated with featureCounts from the subread package v1.5.1 for genes annotated by ensembl Mus_musculus.GRCm38.102, counting the primary alignment in the case of multimapped reads (93). Raw counts were normalized by voom (94). Genes were included if at least half of the samples had an expression of 2 CPM. Heatmaps were plotted with R package ComplexHeatmap. DESeq2 rlog transformation was used to process count data for PCA plotting. Functional enrichment analysis was performed with Ingenuity Pathway Analysis to enrich for IPA Canonical pathways and with clusterProfiler to enrich for KEGG and GO terms. We define FDRsig as FDR<0.05 and psig as p<0.05.

Partek Flow was used for differential gene expression analysis using Gene Specific Analysis (GSA) algorithm by applying multiple statistical models to each individual gene in order to account for each gene’s varying response to different experimental factors, and differing data distribution. Differentially expressed genes (DEGs) were identified for genes showing fold change ≥ 1 and *p-value <0.05*. Wikipathway and Gene Ontology non-redundant biological process enrichment analyses of DEGs among mouse WT and CASP4-KO brain samples were performed using WebGestalt with mouse genome as a reference set (95).

UMAP plots for expression of CASP4 and CXCR3 were generated, as previously described (31), to show expression levels of indicated genes using snRNA-seq datasets from female mouse brain tissues of young and aged animals (GSE207848).

### Statistical analysis

All figures display mean ± standard error of the mean (SEM) from at least three independent experiments as indicated in the figure legends. Comparisons between groups were conducted with either two sample t-test, 2-way ANOVA with Tukey’s correction for multiple comparisons if needed, or mixed effects model with Holm’s adjustment for multiple comparisons as indicated depending on the data structure. Adjusted P < 0.05 was considered statistically significant. Data were analyzed using GraphPad Prism 6.0 (t-test and ANOVA) and SAS 9.4 (linear mixed effects models).

## Declarations

### Ethical Approval

De-identified human samples were obtained in agreement with the NIH Neurobiobank or the Discovery Life Science Biobank. Written informed consent for participation was not required for this study in accordance with the national legislation and institutional requirements. The animal study was reviewed and approved by Animal Care and Use Committee (IACUC) of the Ohio State University College of Medicine.

## Competing Interests

The authors have no competing interests to report.

## Author contributions

Conceptualization, HEG, AOA, KPD.; Formal Analysis, KPD; Investigation, All authors; Resources, AOA.; Writing-Original draft, KPD.; Writing-Review & Editing, All authors; Project Administration, AOA, KPD; Supervision, HEG and AOA.; Funding Acquisition, AOA.

## Funding

Studies in the Amer laboratory are supported by AI24121, HL127651, NIH Covid supplement and a Pilot Grant from the OSU Department of Microbial Infection and Immunity. A.B. is supported in part by the C3 training grant. Images presented in this report were generated using the instruments and services at the Campus Microscopy and Imaging Facility, The Ohio State University. This facility is supported in part by grant P30 CA016058, National Cancer Institute, Bethesda, MD. We would also like to acknowledge the Rodent Behavior Core at the Ohio State University. This facility is supported in part by P30 Core grant (NINDS P30NS04578).

## Availability of data and materials

Methylation and RNA-sequencing data sets are available online as referenced in the text.

---

**Supplementary Figure 1.**
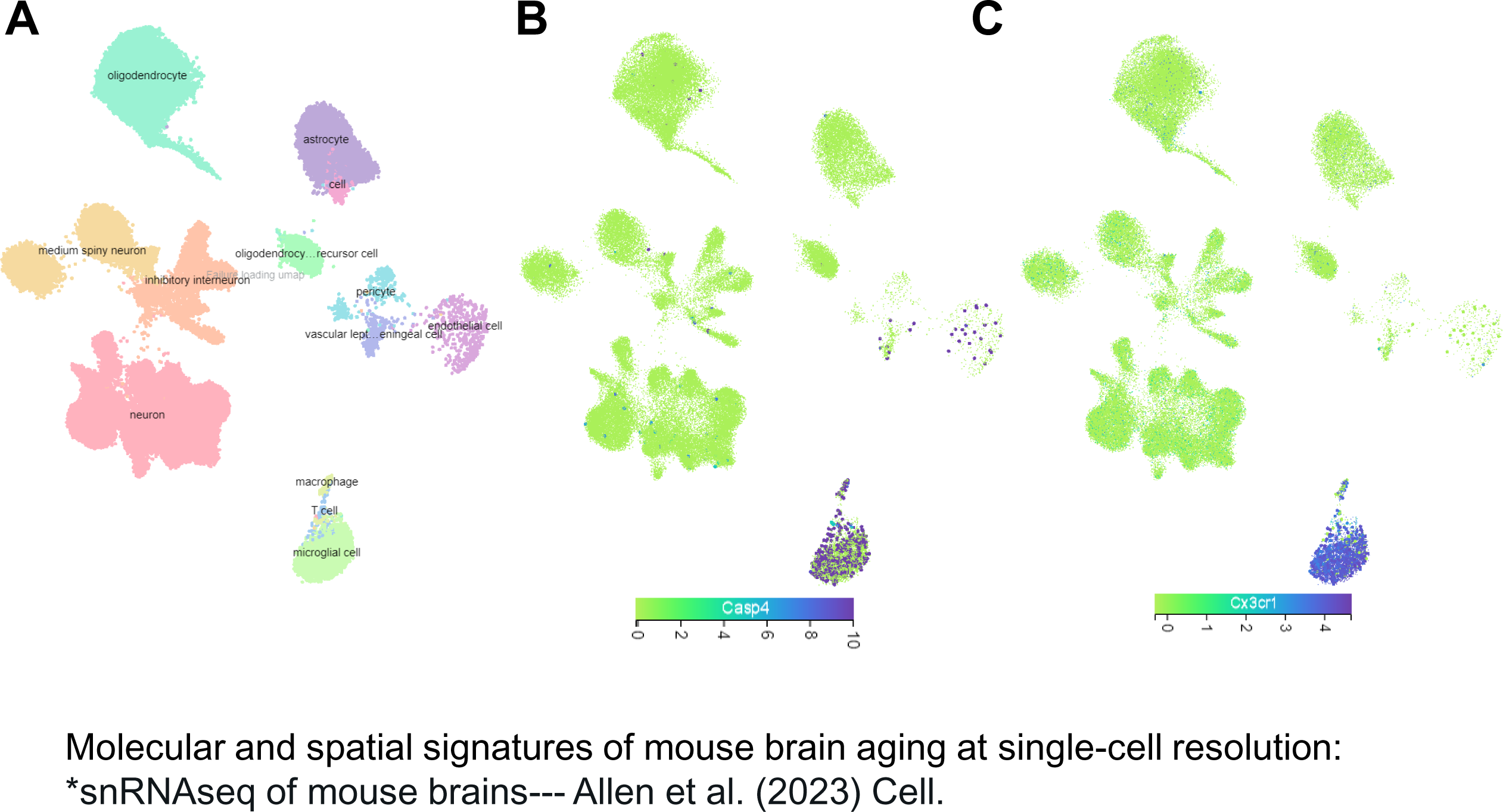
Microglial cells express high levels of Casp4 within mouse brains. A) UMAP visualization of brain cell meta-clusters identified in single-nucleus RNA-sequencing analysis of female mouse brain tissues (GSE207848). UMAP plots showing high RNA expression levels of Casp4 (B) and Cx3cr1 (C) within microglial cells from mouse brain tissues.

**Supplementary Figure 2.**
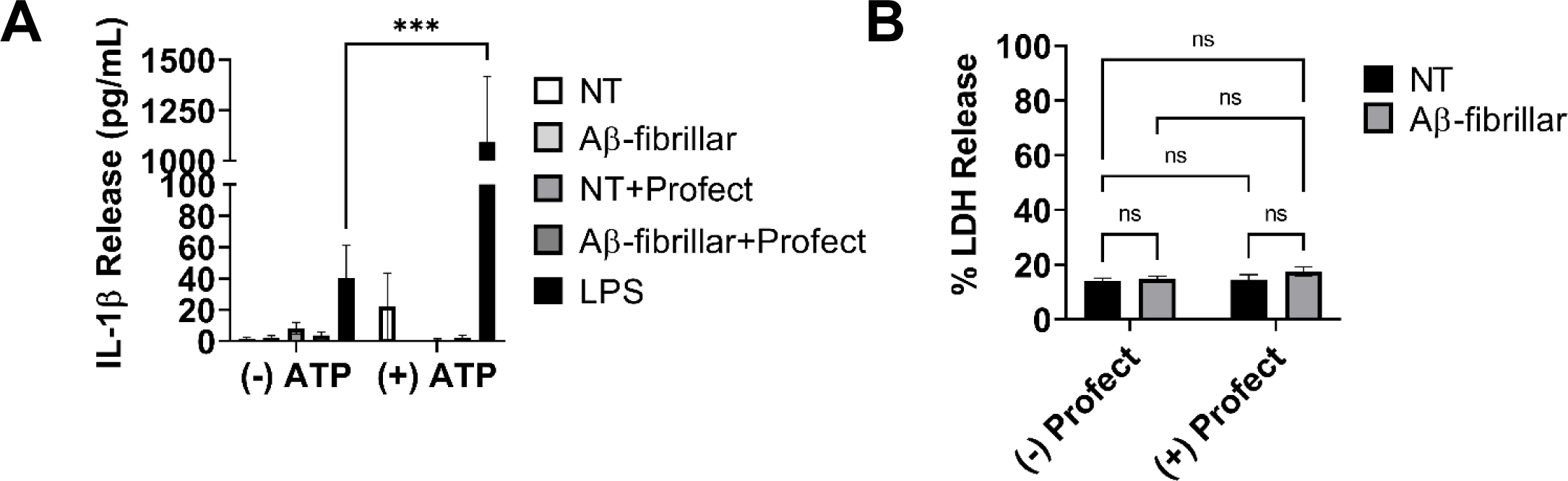
Fibrillar Aβ(1-42) does not prime the inflammasome response or promote cell death. A) IL-1β release from resting macrophages treated for 3 hours with 10µM fibrillar-Aβ (fAβ) or not treated (NT) with and without conjugation to cytosolic delivery reagent Profect or with LPS control followed by 30-minute activation with 5mM ATP (N=4 or N=3 for LPS only). Statistical analysis completed by mixed effects analysis with Tukey’s multiple comparisons test. B) Cell death measured by % LDH release (relative to high control) from LPS-primed mouse macrophages treated for 3 hours with 10µM fibrillar-Aβ with and without Profect, Statistical analysis completed by 2way ANOVA Tukey’s multiple comparisons test (N=5)

**Supplementary Figure 3.**
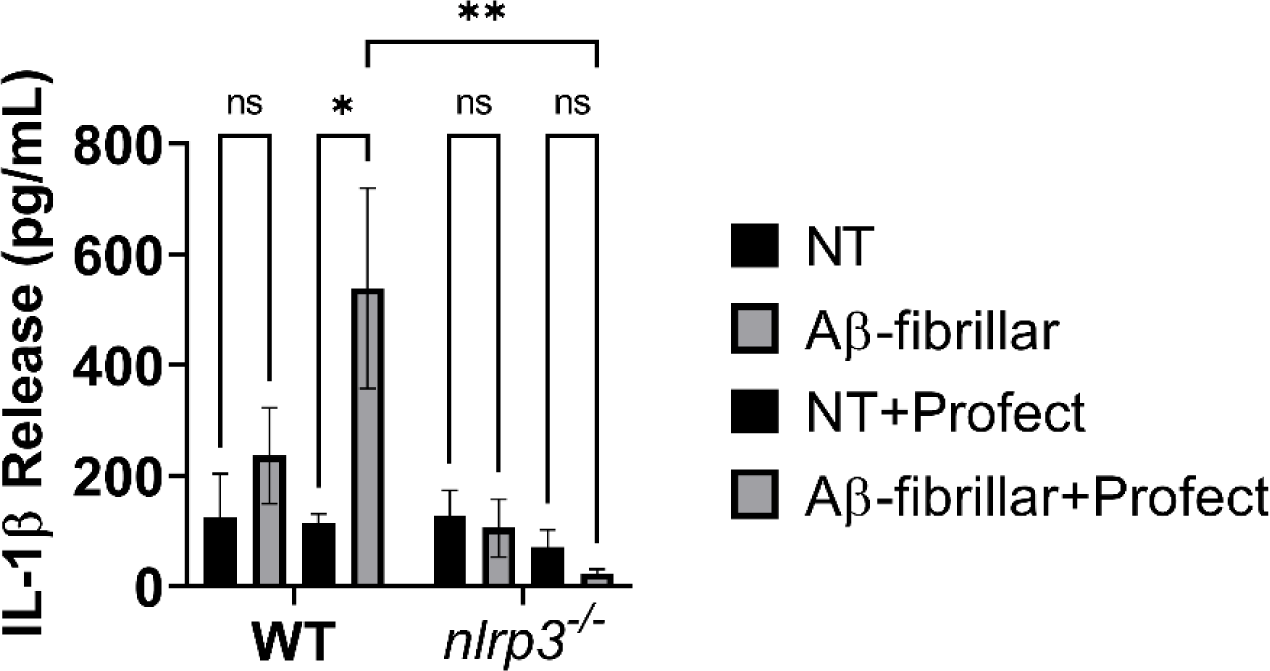
The NLRP3 inflammasome promotes IL-1β release in response to Profect-conjugated-Fibrillar Aβ(1-42). IL-1β release from LPS-primed macrophages from wild-type (WT) and *nlrp3^-/-^* mice treated for 3 hours with 10µM fibrillar-Aβ or not treated (NT) with and without conjugation to Profect (N=3). Statistical analysis completed by 2way ANOVA Tukey’s multiple comparisons test. For simplicity, graph does not display p-values for all comparisons. *P ≤ 0.05, **P ≤ 0.01

**Supplementary Table 1.** Overlap of gene lists with FDR<0.05 when comparing male vs. female mice in a single group.

| Strain | FDRsig | Elements |
| --- | --- | --- |
| 5xFAD,<br>5xFAD/Casp4<sup>-/-</sup> | 5 | Xist Eif2s3y Uty Ddx3y<br>Kdm5d |
| 5xFAD | 1 | Ptgds |

**Supplementary Table 2.** The patient demographic and clinical characteristics of the human samples used in the study. Samples used for methylation analysis are marked in the DMR column. Y: years, BA: Brodmann’s area, h:hours, M:male, F:female, W:White, H:Hispanic, AA:African American, CASP4 DMR: CASP4 differentially methylated region.

| Brain Endowment Bank Tissue order #899 (Temporal Pole (BA38)) |  |  |  |  |  |  |
| --- | --- | --- | --- | --- | --- | --- |
| Non-dementia Brain ID | Label | Sex | Age (y) | Race | Autolysis (h) | CASP4 DMR analysis |
| Hct15HAL_001_18_BA38 | N2 | M | 61 | H | 12.5 | X |
| HctZD_001_18_BA38 | N3 | F | 62 | W | 18.9 | X |
| HctZZC_001_18_BA38 | N4 | F | 82 | W | 14.2 |  |
| HBDE_001_18_BA38 | N5 | M | 83 | H | 29.55 | X |
| HctZZT_001_18_BA38 | N6 | M | 85 | W | 15.5 |  |
| HctYH_001_18_BA38 | N7 | F | 71 | H | 16.4 |  |
| HctYB_001_18_BA38 | N8 | F | 60 | AA | 27.35 |  |
| HctZP_001_18_BA38 | N9 | M | 77 | W | 14.72 |  |
| HctZA_001_18_BA38 | N10 | F | 79 | W | 17.8 |  |
| HctZR_001_18_BA38 | N12 | M | 76 | H | 27.87 |  |
| HctZL_001_18_BA38 | N13 | F | 65 | H | 27 |  |
| Hct15HBC_001_18_BA38 | N15 | M | 83 | H | 25 |  |
| Alzheimer's disease Brain ID | Label | Sex | Age (y) | Race | Autolysis (h) | CASP4 DMR analysis |
| HBDM_001_18_BA38 | AD1 | M | 60 | W | 36.5 | X |
| HBIH_001_18_BA38 | AD2 | F | 62 | W | 3.66 | X |
| HBIQ_001_18_BA38 | AD3 | F | 66 | W | 17.66 |  |
| HBDA_001_18_BA38 | AD4 | M | 80 | W | 22.13 | X |
| HBLZ_001_18_BA38 | AD5 | F | 81 | W | 11.2 |  |
| HBIB_001_18_BA38 | AD6 | M | 82 | W | 11.75 |  |
| HBGU_001_18_BA38 | AD7 | F | 74 | W | 26.5 |  |
| HBFQ_001_18_BA38 | AD8 | M | 77 | W | 18 |  |
| HBEK_001_18_BA38 | AD9 | F | 77 | W | 25.3 |  |
| HBPK_001_18_BA38 | AD10 | M | 78 | W | 24 |  |
| HBFF_001_18_BA38 | AD12 | M | 85 | W | 17.6 |  |
| HBIG_001_18_BA38 | AD13 | F | 60 | W | 7.91 |  |

**Supplementary Data 1 – Filtered DMR list from RRBS**

**Supplementary Data 2 – RNA sequencing, AD/Casp4 KO vs WT**

## Notes

### Competing Interest Statement

The authors have declared no competing interest.

